# Sample preparation optimisation for array tomography volume EM of entire organisms: a case study for marine larvae

**DOI:** 10.64898/2026.10.06.756938

**Authors:** Kata Szabó, Alexandra Kerbl, Karel Mocaer, Angelika Svetlove, Alexandra Pacureanu, Gáspár Jékely

## Abstract

Serial-sectioning electron microscopy allows the exploration of the 3D ultrastructure of cells, tissues or whole organisms. Sample preparation is a critical step for this technique, which determines the quality of downstream imaging and analysis. Here we optimize a high-pressure freezing sample-preparation workflow for larvae of the marine annelid *Platynereis dumerilii.* We tested several fixation and staining methods and screened a large number of specimens by synchrotron X-ray imaging to evaluate specimen integrity and contrast. We further evaluated selected blocks by array tomography SEM to select the best performing protocols. Our protocols and robust workflow could be adapted to similar marine planktonic organisms including ciliated invertebrate larvae.

## Introduction

In recent years, the field of volume electron microscopy (vEM) has rapidly evolved with the rise of new techniques to explore the sub-cellular organization of cells, tissues or whole organisms in 3 dimensions (3D) [1]. In vEM, four approaches are most commonly used, each coming with its own benefits and limitations. Serial block-face scanning electron microscopy (SBF-SEM) [2] uses a knife to subsequently remove a thin layer of specimen from a block between en-block SEM imaging steps. This method enables the imaging of large volumes (up to several hundred µm in each dimension) but is destructive and requires strongly stained specimens. Focused ion beam scanning electron microscopy (FIB-SEM) [3] is generally a method of choice to image smaller regions (below a hundred µm in x, y and z). FIB-SEM has the benefit of acquiring volumes at isotropic resolution, however, it is also destructive. The enhanced version of FIB-SEM (eFIB-SEM) [4] allows the imaging of larger samples (up to 10^6^ µm^3^), but is technically highly demanding and less accessible. Array tomography (AT) [5], [6] allows to image larger regions (up to several hundred µm in each dimension with single-beam SEM) while preserving the sections thus permitting re-imaging specific areas at higher resolution. AT also enables the introduction of an additional on-section contrasting step. This technique is also more widely accessible than FIB-SEM or SBF-SEM as it only requires a sample-preparation workflow and an SEM. Lastly, serial-section TEM uses serial ultrathin sectioning on slotted grids or tape and images sections in transmission mode. TEM allows faster imaging and gives better image contrast, also potentially enabling high-resolution (sub-nm) imaging [7], [8]. The downsides include the use of non-rigid substrates and the challenges in automation [1], although techniques to counteract these disadvantages are being developed [9].

Sample preparation is a critical step in each of these methodologies. This is particularly the case when fixing entire organisms or environmental samples rather than conventional tissue samples (e.g. brain sections or histological samples). The challenges include the presence of a cuticle, ciliary bands, mucus or various appendages [10]. Different types of organisms often require individually optimised protocols. For example, for the miniature wasp *Megaphragma amalphitanum* a special chemical fixation protocol has been developed to allow fixation and staining through a thick chitin cuticle [11].

Here our aim was to develop a protocol suitable for array tomography for small whole organisms such as ciliated marine larvae. We focused on larvae of the marine annelid *Platynereis dumerilii.* Previous vEM work with this organism was performed using either ssTEM or SBF-SEM. A chemical fixation protocol gave limited preservation in the neuropil of the 6-day-old larva analysed by SBF-SEM, precluding synapse detection [12]. The 3-day-old ssTEM volumes were collected from specimens prepared by high-pressure freezing (HPF) and enabled synaptic connectomics [13], [14]. However, specimens show lower contrast in SEM and the original workflow did not include an X-ray-based pre-screening step and the specimens showed smaller gaps and some tissue damage.

Here we focused on optimising procedures for *Platynereis* larvae using high-pressure freezing (HPF), a method known to give the best quality of ultrastructural preservation [15], [16], [17]. We evaluated all samples by synchrotron X-ray [18] imaging and serial-section SEM to select intact specimens and the best protocol.

## Materials and methods

The catalogue numbers of all reagents, consumables and equipment is included in the supplementary section (Supplementary Table 1).

### Culture of animals

*Platynereis dumerilii* [19] were cultured using the method originally established by Hauenschild (1951) [20] and further developed by Fischer and Dorresteijn (2004) [21]. In our cultures, we also implemented more recent improvements suggested by Kuehn *et al.* (2019) [22], Legras *et al.* (2023 [23]), and Hird et al. (2024) [24].

### Fixation, staining and embedding of the samples

#### High pressure freezing

We loaded 6-day-old *Platynereis dumerilii* larvae in the 200 µm side of a 3-mm type A gold/copper or aluminum carriers and covered them with the flat side of a 3-mm type-B aluminum carrier. The carriers were previously coated in hexadecene. We then high-pressure froze the specimen using an BAL-TEC HPM 010 high-pressure freezer.

#### Freeze substitution

Following high-pressure freezing, the vitrified samples were freeze-substituted using an AFS2 freeze substitution unit. We tested different protocols, each of them are detailed below and summarised in Supplementary tables 2-5. We embedded all the samples in LX112 resin (hard mix) from Ladd Research.

We tested two different methods to process the carriers. The first method used screw-cap cryo-tubes containing the freeze-substitution mix, with one carrier per cryo-tube. Every exchange was done in the cryo-tubes, until resin embedding where we transferred the specimens into convex glass dishes. Ultimately we transferred the samples to molds using toothpicks. The second method used the buckets for freeze substitution [15] to load the A carriers containing the specimen facing up.

The protocols in table form are in Supplementary tables 2-5.

##### Protocol A – Adapted from EMCF Nice

The cryopreserved samples were placed in the freeze substitution machine for 37 h at -90 ℃ in the freeze substitution cocktail: 1% osmium tetroxide (OsO_4_) 0.5% uranyl acetate (UA) and 3% water in dry acetone. The temperature then increased to -60 ℃ over 15 h at a rate of 2 ℃/h. The samples stayed at -60 ℃ for 15 h. Another step to -30 ℃ followed over the course of 15 h at a 2 ℃/h rate, we then kept specimens at -30 ° C for 12 h. We then increased the temperature to 0 ℃ over 1 h, at a 30 ℃/h rate. The samples stayed at 0 ℃ for 1 h after which they were cooled down again to -30 ℃ over at a rate of 30 ℃/h. Once the temperature reached -30 ℃, we rinsed the samples with acetone every 15 minutes 5 to 6 times.

After the acetone rinses, we facilitated resin infiltration of the sample through a series of increasing amounts of resin without accelerator mixed with acetone. We used the following resin:acetone ratios (v:v): 30% (2 h from -30 ℃ to -10 ℃, 10 ℃/h), 50% (2 h from -10 ℃ to 10 ℃, 10 ℃/h), 70% (2 h from 10 ℃ to 20 ℃, 5 ℃/h). After the 70% resin exchange step we exchanged the solution to 100% resin without accelerator at room temperature (RT) and let the samples infiltrate overnight, next we exchanged to 100% fresh resin again and left them for an additional 48 h. We then exchanged the resin to 100% resin with the accelerator DMP30 3 times for 3 h each. The samples kept in tubes were then transferred into molds (which were filled with a prepolymerized layer of resin for optimal positioning of the specimen), and let settle overnight. After the overnight settling step samples were kept in a 60 ℃ oven for 48 h.

##### Protocol B – Adapted from unpublished work from Dr Motohide Murate

The freeze-substitution started at -90 ℃, the samples were kept at this temperature for 54 h in 2% OsO_4_, 0.5% UA, 1% glutaraldehyde (GA) in acetone. The temperature was then increased to -20 ℃ over 8 h at a rate of 8.75 ℃/h. The samples were kept at -20 ℃ for 12 h, afterwards the AFS was warmed to 4 ℃ at a 10 ℃/h heating rate. Samples were washed with acetone 4 times every 19 minutes at 4 ℃ followed by an incubation in 1% UA (in acetone) for 14.5 h, staying at 4 ℃. The machine was then set to 20 ℃ over 30 min (32 ℃/h rate), we followed this by 5 times 10 minute acetone washes. Samples were removed from the AFS machine and the subsequent steps were carried out at room temperature. For resin infiltration, we used a series of resin:acetone mixtures (25%, 50%, 75%, 100%) for 2 h each, the mix did not contain accelerator. The samples stayed in 100% resin overnight. The next morning the resin was changed again, in which the samples stayed for 48 h. After that the resin was changed again 3 times, each step taking 4 h, this time however the accelerator was added to the resin. After the last change, some samples were transferred into molds on a layer of pre-polimerized resin, the rest of the samples were left in the carriers. The samples were left to settle overnight, and were transferred the next morning into a 60 ℃ oven for at least 48 h.

##### Protocol C – Adapted from Stigloher et al, 2011 [25] and Gallet et al, 2024 [15]

The samples were placed in the freeze-substitution machine for 28 h at -90 ℃ in 0.1% tannic acid (TA) and 0.5% GA in acetone followed by acetone rinses every 15-20 min for 1.5 h. A new freeze-substitution mix of 2% OsO_4_ in acetone is added to the samples at -90°C, where they stay for 29.5 h. Then the temperature is increased to -20 ℃ at a 5 ℃/h rate for 14 h. The samples stayed at -20 ℃ for 16 h, the temperature was increased again to 4 ℃ at a 6 ℃/h rate for 4 h. When the temperature reached 4 ℃, samples were rinsed with acetone every 15-20 minutes for 1.5 h. A 25% resin:acetone (without accelerator) mix was added to the samples, while the temperature was increased to 20 ℃ at 8 ℃/h for 2 h. An increasing resin series (50%, 75% then 100%) was added to the samples with 2 h of incubation in each solution. After the last step, the samples were transferred out of the AFS to room temperature, where they stayed overnight. Samples could be transferred out of the AFS sooner, as the temperature reaches 20 degrees. The next morning another resin change was done in which the samples were left for 48 h. This was followed by three incubations in resin with accelerator (0.014 g per 1g of resin) for 4 h each. After the last step, part of the samples were transferred to molds with pre-polimerized resin, while the rest were left in the carriers, and all the samples were left to settle overnight at room temperature. This was followed by a polymerization step in a 60 ℃ oven for at least 48 h.

##### Protocol D – Adapted from Tsang et al. 2018 [26]

The freeze substitution solution 1 (0.2% GA, 0.1% UA, 2% MeOH, 1% H_2_O) was cooled down to -120 °C in the freeze-substitution bucket before carriers were transferred into the wells. Then we increased the temperature at a heating rate of 7.5 ℃/h for 4 h from -120 ℃ to -90 ℃. The machine stayed at -90 ℃ for 53 h, after this the temperature was increased to -60 ℃ in 15 hours, at a 2 ℃/h rate. Samples stayed at -60 ℃ for 15 h followed by a heating step to -30 ℃ for 15 h at a 2 ℃/h rate. The AFS was kept at -30 ℃ for 14 h, after this time we washed the samples three times with 0.2% GA, 1% water in acetone, at 20 min intervals. A quick heating step from -30 ℃ to 0 ℃ followed, this took 5 min at a 360℃/h rate, while the samples stayed in the washing solution.

Samples were transferred to ice, where 10 re-hydration steps (20 minutes each) followed, with the following solutions (RS1 – RS9). RS1: 5% water, 0.2% GA in acetone, RS2: 10% water, 0.2% GA in acetone, RS3: 20% water, 0.2% GA in acetone, RS4: 30% water, 0.2% GA in acetone, RS5: 50% 0.1M HEPES (buffered to pH 7.4), 0.2% GA in acetone, RS6: 70% 0.1 M HEPES (buffered to pH 7.4), 0.2% GA in acetone, RS7: 0.1 M HEPES (pH 7.4), RS8: 0.15 M sodium cacodylate with 100 mM glycine (pH 8.0), RS9: 0.15 M sodium cacodylate (pH 7.4). We then kept half of the samples in carriers, while we took out the other half to convex-glass dishes to later be embedded individually.

While staying on ice, we stained the samples with a round of heavy-metal-containing solution. The samples were kept in the mixture (2% OsO_4_/1.5% potassium ferrocyanide/2 mM CaCl_2_ in 0.15 M sodium cacodylate, pH 7.4) for 1 h, after which we washed them five times for 5 minutes. We removed the samples from the ice, to continue the heavy metal staining for 30 min with 0.5% thiocarbohydrazide (TCH) in water (adjusted to pH 7.0, filtered through a syringe). Another five times 5 min washes with water followed. We then did a third round of heavy metal staining with 2% of OsO_4_ in water (pH 7.0), followed by five 5 min washes in water. We did an overnight staining step in 2% aqueous UA (pH 4.4, filtered through a syringe), then 5 more times 5 min washes with water.

Following the heavy metal staining, samples were dehydrated in an EtOH series (70%, 90%, and 2 times 100%) for 10 min each. Two 10 min washes with 100% acetone followed to prepare the samples for optimal infiltration with the embedding medium. This step was necessary due to the strongly hydrophobic nature of LX112. The samples were infiltrated with LX112 resin without accelerator in a resin:acetone series consisting of 2 h in 25%, 50%, and 75% resin. Samples were then transferred to 100% resin, in which they stayed overnight. The following day, 3 exchanges of 100% fresh resin with accelerator were done, each incubation lasting 4 h followed by an additional overnight incubation. For the last change we took out the samples of the carriers and transferred them into molds that had a bottom layer of pre-polimerized resin. Following the overnight incubation the samples were transferred into a 60 ℃ oven for resin polymerisation for at least 48 h.

##### Protocol E and F

We treated samples with 4% paraformaldehyde solution (PFA) for one hour prior to HPF. Then we freeze-substituted them following either protocol C for protocol E and D for F.

### Resin preparation and embedding

We embedded samples, in either molds or freeze-substitution buckets (Figure 1). Good quality samples embedded using buckets were cut out with a razor blade and re-embedded after X-ray scanning at the Hamburg P14 beamline.

**Figure 1:**
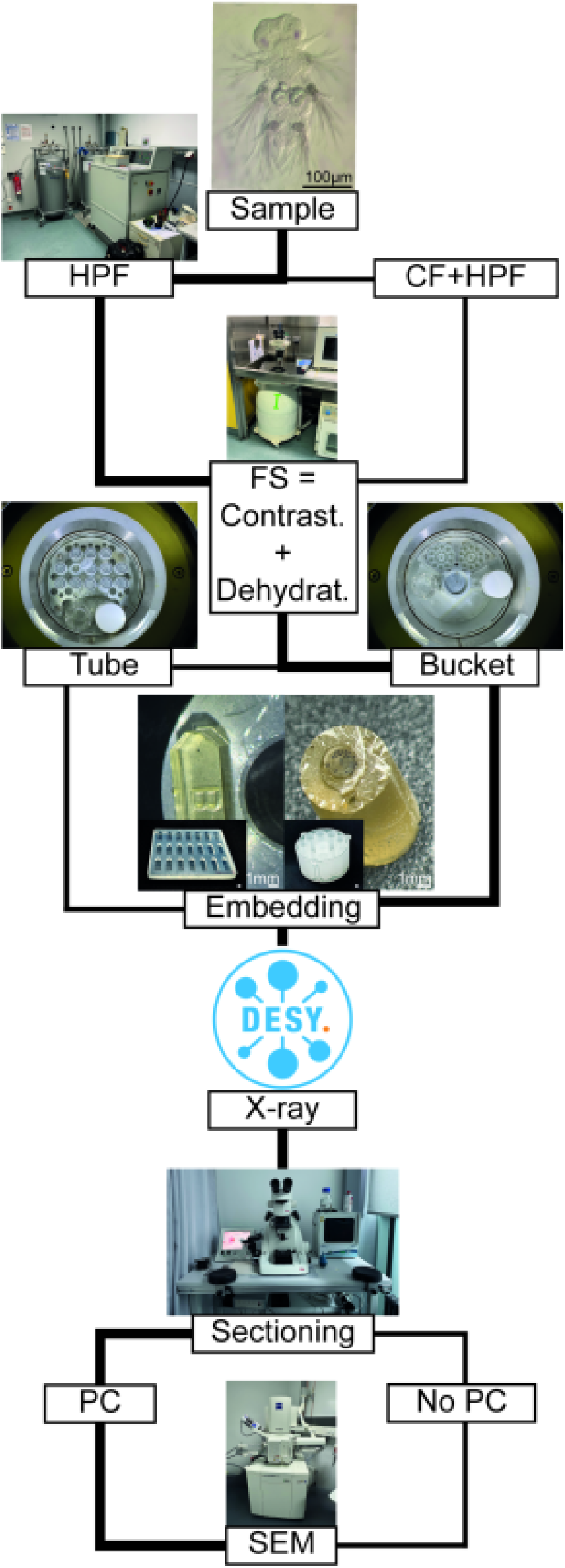
Schematic of different steps optimized for the workflow for whole-body volume EM and connectomics using array tomography. Abbreviations: FS -freeze-substitution, HPF -high-pressure freezing, HPF+CF -high-pressure freezing following mild chemical fixation, PC -post contrasting with 3% lead citrate. We indicated the workflow which we found most beneficial for sample preservation in bold lines.

We used the epoxy-based resin LX112, mixed the resin according to the manufacturer’s instructions to hard (3:7 w:w) relative hardness. We prioritized LX112 resin as it is considered more suitable for array tomography, as it gives less compression artefacts as opposed to EPON [29], [30].

### X-ray

#### Hamburg – P14

We trimmed the blocks containing a single specimen using a 90° diamond trimming knife . We did it so the sample would be located in a small cuboid shape with a few 100 micrometers of resin surrounding the larva, with the bottom still attached to the original resin block. As for the samples embedded with carriers we cut them out of the buckets and carefully removed the carriers from them using razorblades. The blocks were then mounted on 3D printed stubs to be the correct height. Each block was imaged at the Deutsches Elektronen-Synchrotron (DESY), Hamburg, at the P14 beamline [31]. Specimens were imaged at four propagation distances of 73, 77, 83 and 92 mm at 18 and 23.5 keV with 650 nm and 325 nm voxel sizes. Trimmed blocks were imaged using a standard in-centre acquisition with 1800 projections, the HPF blocks were imaged off-centre with 3600 projections and an overlap of ∼250 voxels. Phase retrieval was performed using 0.1 beta/delta ratio. TOMO-CTF software [32] was used for phase retrieval and tomographic reconstruction. The acquired data was uploaded to Webknossos [33], for storage and visualization.

In order to assess the contrast-to-noise ratio (CNR) of each protocol, we used the biological image-analysis platform FIJI [34]. We imported 10 randomly selected samples from each sample preparation protocol from Webknossos to Fiji as Tiff stacks. For each sample we chose 3 ROIs in the head of the animals to assess the contrast and 3 ROIs in the resin as background. From this data we calculated the CNR using the following equation:

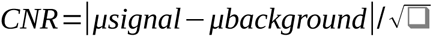

#### Grenoble – ID16A

For the ID16A scans, samples had to be re-embedded after scanning them at the P14 beamline. Some of the samples were re-trimmed so that they could be safely cut off from the block. We re-embedded the samples in a 20 μl pipette tip in EPON to ensure that samples had a uniformly smooth surface. This mounting method limits undesirable X-ray scatter and enables higher resolution imaging.

The whole sample was then imaged at a 100 and 55 nm voxel size, and the head was imaged at 20 nm resolution. For the tomography, several projections with equal angular spacing between sample rotations of 0° and 180° were recorded. The tomograms were reconstructed using a ‘rigid’ algorithm.

### Sectioning, post-staining and imaging

We sectioned the samples to 40 or 50 nm thickness using a UC7 ultramicrotome and a diamond knife, then collected the sections on ITO-coated glass slides, which had been glow discharged following the manufacturer program . We transferred the sections to an oven (to dry at either 37 or 50 ℃ for at least 1 hour. Later we post-stained some of the sections using 3% lead citrate for 5 minutes, then rinsed the sections with water and air-dried them. We then imaged the sections on a Zeiss GeminiSEM 500 or Zeiss GeminiSEM 360, using the Zeiss Sense BSD detector operating at a working distance of 4.0 mm. We set the EHT to 1.50 kV, the aperture to 30 µm, and imaged at high current (220 pA (0.22 nA / 0.00022 mA)). For images we used a resolution of 2048 x 2048 px, with a pixel dwell time of 25.6 µsec.

All images analysed and presented in this study have been automatically adjusted for brightness and contrast in Fiji.

In order to objectively assess and compare signal-to-noise ratio (SNR) and contrast of each protocol also on the EM images, we employed the FIB-SEM-plugin developed by Shtengel et al. [35] for Fiji. This tool is primarily designed to analyse signal-to-noise ratio, evaluate contrast, and assess resolution in single images in FIB-SEM datasets. Regardless of the different nature of the samples, we applied the tool to individual images from the datasets in the approximate center of the animals under the premise that all datasets were acquired with similar settings. Using default settings, we calculated SNR and wherever possible also contrast, yet the latter was often obstructed by the small difference between the data and the surrounding resin.

## Results

We optimized a sample preparation workflow (Figure 1) for *Platynereis dumerilii* larvae for AT vEM. The larvae range from 200 to 350 μm in length and are composed of ∼10,000 cells. Several different parameters were optimized, starting from the fixation of the samples. We tried four HPF protocols and compared it to mixed chemical-HPF protocols. The samples were processed by different freeze-substitution protocols varying in staining reagents and timings. To test sample integrity and staining uptake, we carried out X-ray imaging on the P14 beamline at DESY in Hamburg, Germany. We also assessed the quality of the preparations by imaging ultrathin sections with SEM. A few specimens were also tested with high-resolution X-ray imaging at the ESRF ID16A beamline.

### Optimisation of specimen handling during high-pressure freezing to reduce breakage within tissue

HPF is known to introduce cracks in the samples precluding the collection of comprehensive vEM data. We scanned samples at the P14 beamline at DESY in Hamburg to check tissue integrity. In most specimens prepared by HPF, cracks were apparent. Our first goal was to minimise this undesired in-tissue cracking to enable the collection of intact volumes. We tried various cryo-protectants, different carriers, and different methods of specimen handling (Table 3) and density.

The scanned larvae were categorized based on severity as A) no detrimental cracks, sample suitable for whole-body reconstruction B) small crack in an important region of the sample (e.g., head), C) big crack that would preclude volumetric analysis such as connectomic reconstruction, D) missing parts of tissue or poorly preserved sample (E) (Table 1, Figure 2A-2E).

**Figure 2:**
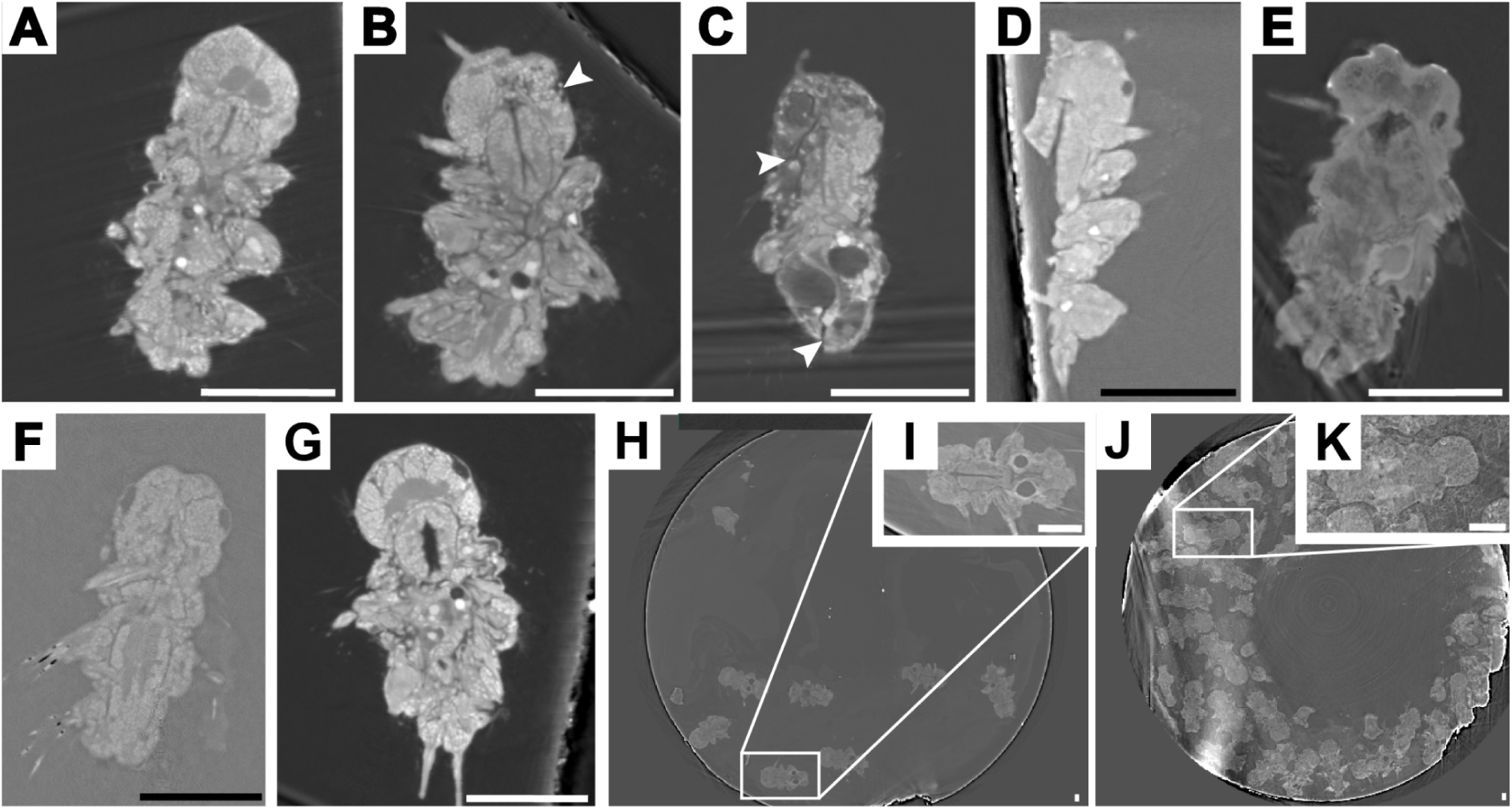
Evaluation of samples by synchrotron X-ray imaging revealing different extent of tissue damage (arrowheads indicating cracks). (A) Intact sample, (B) small crack in anatomically important location (head), (C) big crack or multiple cracks in the body, (D) missing tissue, (E) intact samples that look abnormal. The amount of heavy metal stain that a sample took up can also be assessed from the X-ray images. (F) A well stained sample, (G) a less stained sample. We can also assess the quality and identify good quality larvae (I) in samples where the larvae were embedded in buckets (H). However, it is not possible if the number of animals embedded together is too high, as it leads to higher diffraction of the X- rays, lowering the resolution. Scale bars 100 μm.

**Table 1:** Quantification of damage in the samples based on X-ray images. Damage severity was assessed by eye by analysing the entire image stack of each block, which included either individual or several samples. Not all individuals could be scored in all blocks due to overlap, edge effects or occlusion by the cryoprotectant.

| Sample quality | Tube | Bucket |
| --- | --- | --- |
| Intact | 35 larvae (33 %) | 76 (44 %) |
| Small crack in anatomically important location (head) | 31 (29 %) | 31 (18 %) |
| Big crack | 19 (18 %) | 31 (18 %) |
| Missing tissue | 12 (11 %) | 33 (19 %) |
| Intact, but poorly preserved | 9 (8 %) | 0 (0 %) |
| Total | 106 larvae from around | 171 larvae from 24 |
|  | 30 planchets fixed in 13 substitution rounds | planchets fixed in 7 substitution rounds |

First, we tried various cryoprotectants, including polyvinylpyrrolidone (PVP) and yeast paste. For samples frozen with PVP, we did not notice a reduction in cracks. We also tried using yeast or varying the number of animals in the same carrier. In case the samples were freeze-substituted in a free-floating manner (outside of the carriers), we did not notice a reduction in the number of damaged samples. When we used the buckets to freeze-subtitute the samples within the carriers, however using both yeast or a larger number of animals caused the same problem: the X-ray images lacked enough resolution to determine the quality of the samples. This was likely due to the scatter or absorption of the X-ray by the high density of well stained specimens or yeast.

Second, we froze a varying number of animals in the same carrier. When over 25-30 specimens were embedded in the same carrier, the crowding resulted in a drop in the resolution in the X-ray scans. This made it difficult to distinguish and cut out individual specimens for high-resolution X-ray imaging or AT. However, we found that freezing between 10-15 larvae together works well both for X-ray and for the subsequent isolation of individual specimens (we did not notice an impact on freeze damage).

Third, we tried different methods of specimen handling during freeze-substitution. We tried both screw-cap cryo-tubes and buckets (Figure 1 and Methods). While the specimens generally moved out of the A carriers using the first method (screw-cap cryo-tubes), they generally stayed in position using the second method (buckets), avoiding the loss of small samples (30-100 µm) which can be complicated to locate during exchange steps.

Finally, we tested both aluminum- and gold-coated copper carriers. We noticed that the samples were better stained following HPF with the gold carriers. Furthermore, sometimes the staining agents would precipitate when in the aluminum carriers, a problem we did not encounter with the gold-coated ones.

Overall, we found that freezing 10-15 animals without cryo-protectant in gold-coated copper carriers and handling them in the freeze-substitution buckets yielded samples optimal for X- ray imaging and showing the least number of cracks (summarised in Table 1).

Out of the more than 250 individually embedded larvae, 30-40% showed no or only minimal in-tissue breakage. For the samples embedded in the bucket, from around 50 scanned blocks 40% of the specimens were classified as adequate for re-embedding and sectioning.

These results highlight the critical importance of X-ray scanning of large numbers of specimens before proceeding to large vEM reconstruction requiring a fully intact specimen.

### High-resolution X-ray imaging of high-pressure-frozen specimens

To further assess specimen contrast and preservation at higher resolution, we turned to high-resolution X-ray imaging at the European Synchrotron Radiation Facility (ESRF) in Grenoble, France. We selected two good quality samples from the samples scanned at the P14 beamline in Hamburg and re-embedded them for X-ray imaging at the ESRF. The first sample (prepared with protocol A) showed lower contrast and was difficult to locate in the block. The second sample (prepared with protocol C) had higher contrast and was used for all subsequent imaging steps.

We recorded three datasets, including one at a resolution of 100 nm (Figure 5A) containing the whole larval body, one at 55 nm (Figure 5B) also containing the whole body and one at 20 nm (Figure 5C) focusing on the head. Following reconstructions, we found that our 55 nm sample showed good contrast in the nuclei, however lacked contrast in the membranes.

Obtaining better membrane contrast will require further optimisations of the staining protocol. We also noticed that the block started to show beam damage when imaged at 20 nm resolution (Supplementary Figure 2). This is a known problem and mitigating it would require embedding the samples in a beam-resistant resin such as TGPAP [36].

### Assessment of various staining protocols by X-ray imaging and serial-section SEM

To identify the best method for staining and preservation, we tried various fixation and freeze-substitution protocols. In protocols A-D we combined HPF with freeze-substitution in various solutions. In protocols E and F we introduce a chemical fixation step before HPF and freeze-substitution. The protocols are summarized in Table 2 and detailed in Supplementary tables 2-5.

**Table 2.** Summary of the different fixation and staining protocols employed in this study. Abbreviations: CF -chemical fixation, EtOH -ethanol, FA - formaldehyde, FS - freeze- substitution, GA - glutaraldehyde, HPF - high-pressure freezing, CF+HPF - mild chemical fixation followed by high-pressure freezing, OS - osmium tetroxide, PC - post-staining with 3% lead citrate, PFA - Para-formaldehyde, TA - tannic acid, TCH - thiocarbohydrazide, UA - uranyl acetate.

| Protocol | Fixation and staining | Resin | Comments |
| --- | --- | --- | --- |
| (A) HPF + FS | 1% OsO<sub>4</sub> + 0.5% UA | LX112 | Well preserved ultrastructure, slightly less staining |
| (B) HPF + FS | 2% OsO<sub>4</sub>, 0.2% UA, 0.1% GA -> 0.2% UA | LX112 | Well preserved ultrastructure and well stained samples |
| (C) HPF + FS | 0.1% TA + 0.5% GA -> 1% OsO<sub>4</sub> | LX112 | Well preserved ultrastructure and well stained samples |
| (D) HPF + FS | 0.15% GA, 0.2% UA -> 4% GA -> GA + HEPES -> OsO<sub>4</sub> + TCH + potassium ferrocyanide | LX112 | High amount of staining uptake, slightly worse preserved ultrastructure |
| (E) CF + HPF + FS | 4% PFA -> 0.1% TA + 0.5% GA -> 1% OsO<sub>4</sub> | LX112 | Slightly worse preserved ultrastructure, well stained samples |
| (F) CF + HPF + FS | 4% PFA -> 0.15% GA, 0.2% UA -> 4% GA -> GA + HEPES -> OsO<sub>4</sub> + TCH + potassium ferrocyanide | LX112 | Slightly worse preserved ultrastructure, well stained samples |

**Table 3:** Number of samples (blocks) per condition and protocol. The total number of individuals could not be determined for each block due specimen crowding or the occlusion by the cryo-protectant in some blocks prepared by the bucket method.

| Protocol | Tube | Bucket |
| --- | --- | --- |
| Protocol A | 17 | 3 |
| Protocol B | 7 | 8 |
| Protocol C | 47 | 39 |
| Protocol D | 17 | 4 |
| Protocol E | 2 | 4 |
| Protocol F | 7 | 4 |

First we compared the contrast-to-noise ratio (CNR) of X-ray images collected at the P14 beamline. Protocol D had the best CNR, while protocols C and E had the lowest (Figure 3), although there was considerable variance in CNR between samples prepared with the same protocol.

**Figure 3:**
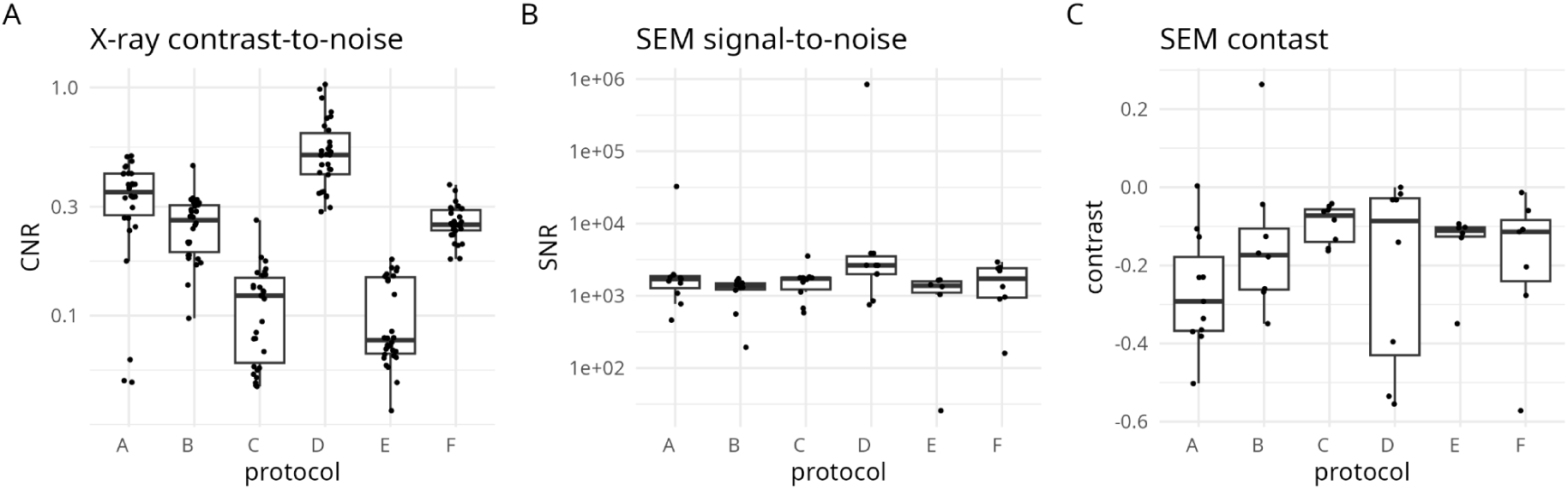
Assessment of contrast of the different protocols. (A) Contrast-to-noise ratio of the different sample preparation protocols based on the X-ray data. (B) Signal-to-noise ratio and (C) contrast of the different sample preparation protocols based on SEM data.

Next, we used serial sectioning and SEM imaging to systematically compare all protocols. We collected ribbons of sections on conductive ITO glass slides and dried them in a 50 ℃ oven. Occasionally we observed lacunae under the sections. When we dried the ribbons at 37 ℃, we did not notice this phenomenon anymore therefore we used 37 ℃ for all samples (we did not investigate the effect of drying temperature systematically).

In initial tests we sectioned samples at both 40 (Figure 6A) and 50 nm (Figure 6B) thickness [10]. The 40 nm sections were more fragile, the membranes showed fainter signal and we sometimes also detected other artefacts on the sections while imaging, possibly from drying or post-staining. The 50 nm sections were easier to manipulate and formed more consistent ribbons. We therefore did all subsequent comparisons with 50 nm sections.

**Figure 4:**
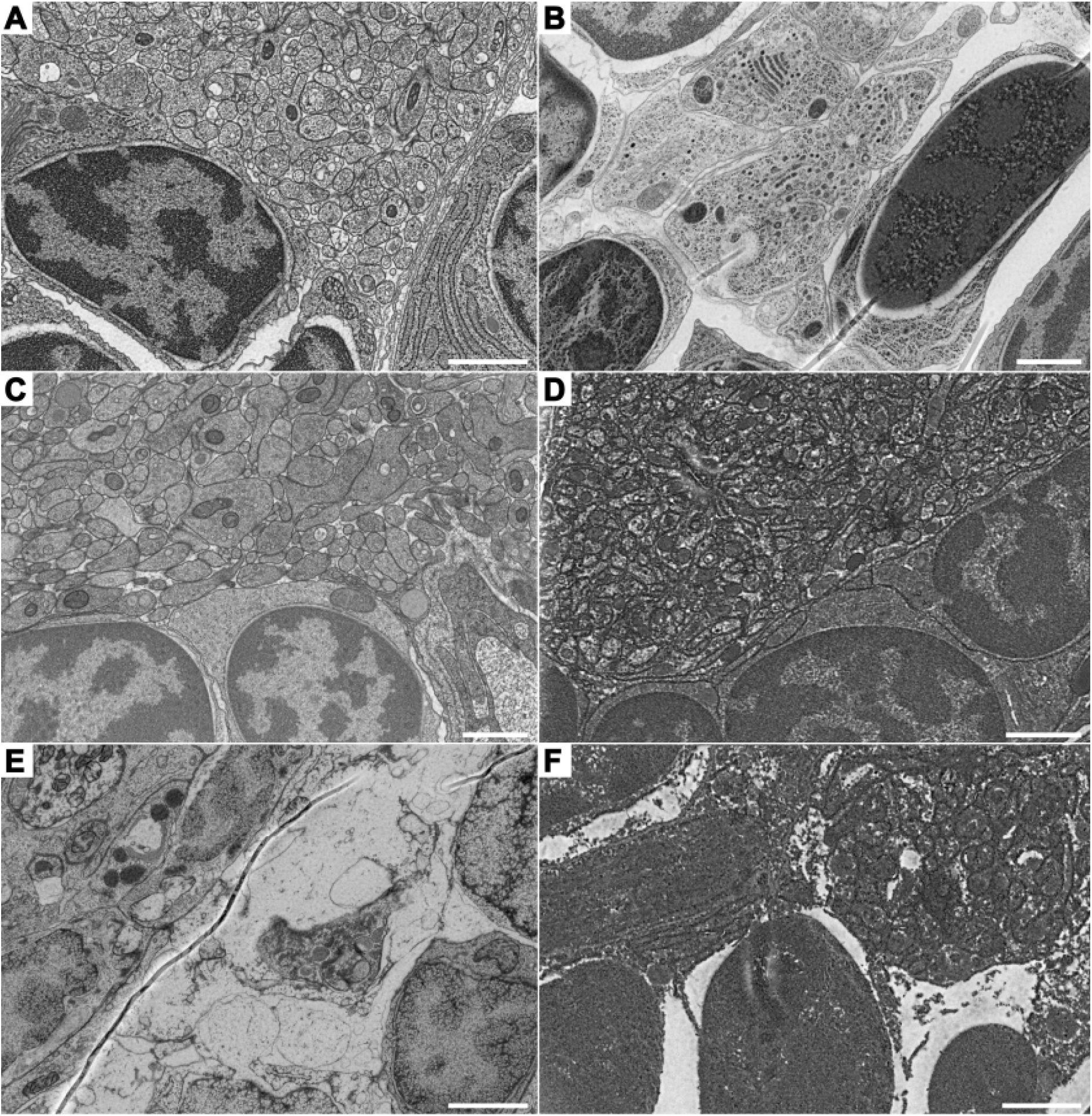
Evaluation of sample preparation protocols by EM imaging, showing similar regions in the head of the 6-day-old *Platynereis*. (A) Protocol A shows good preservation of ultrastructure. (B) Protocol B also shows good preservation of ultrastructure, however some folds can be seen. (C) Protocol C also shows good ultrastructural preservation. (D) Protocol D shows good ultrastructure preservation, however the high contrast makes it harder to distinguish them. (E) Protocol E shows poor ultrastructural preservation, with undulated membranes, poorly preserved organelles and folds are also present. (F) Protocol F similarly to E shows poorly preserved ultrastructure, while also having high contrast, making it difficult to discern details.

**Figure 5:**
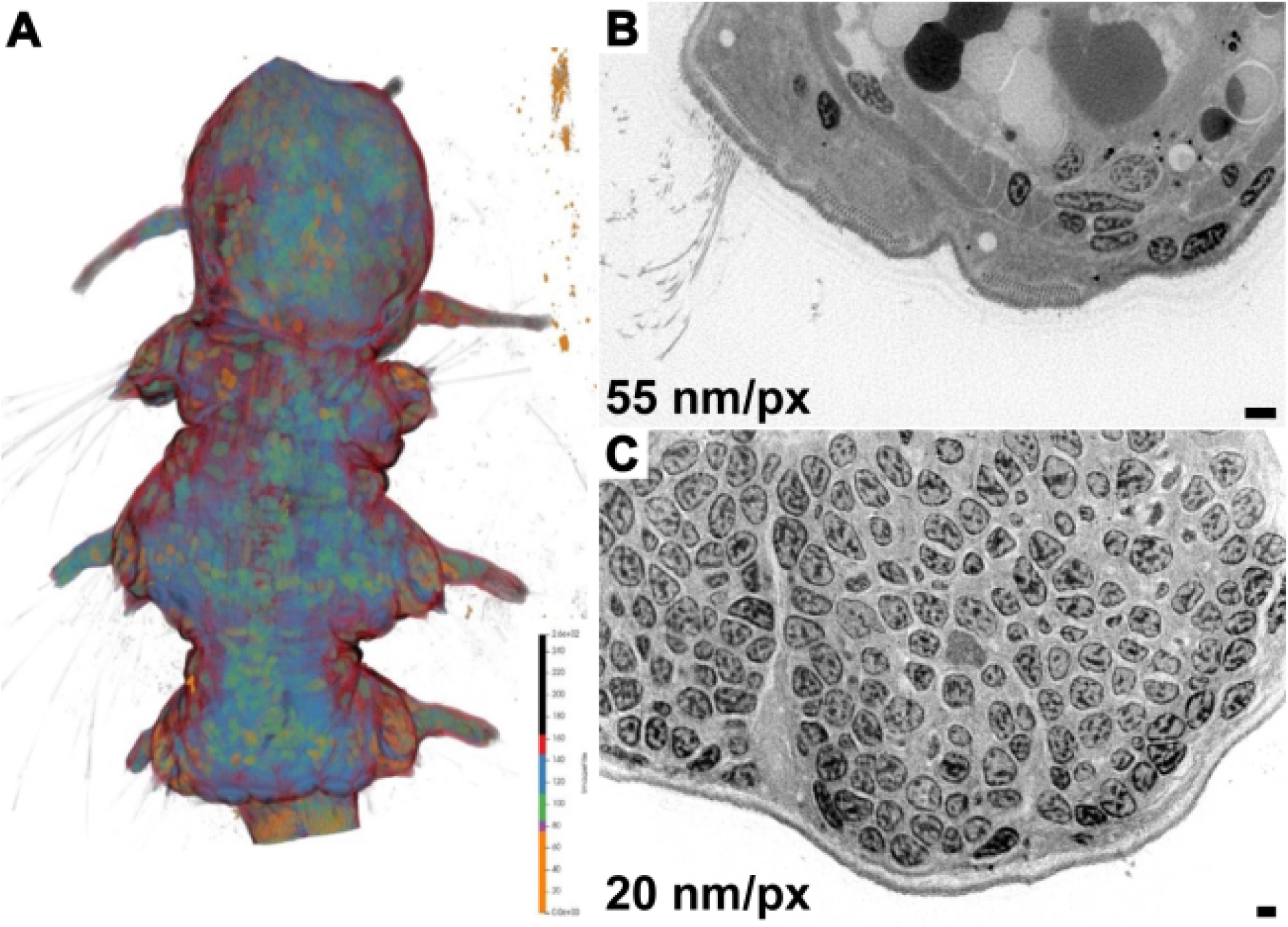
X-ray imaging at ESRF. (A) Morphological rendering of a 6-day-old *Platynereis* prepared with Protocol C and scanned at 100 nm/px. (B) Image plane of the same larva scanned at 55 nm/px or (C) 20 nm/px. Scale bars 1 μm.

**Figure 6:**
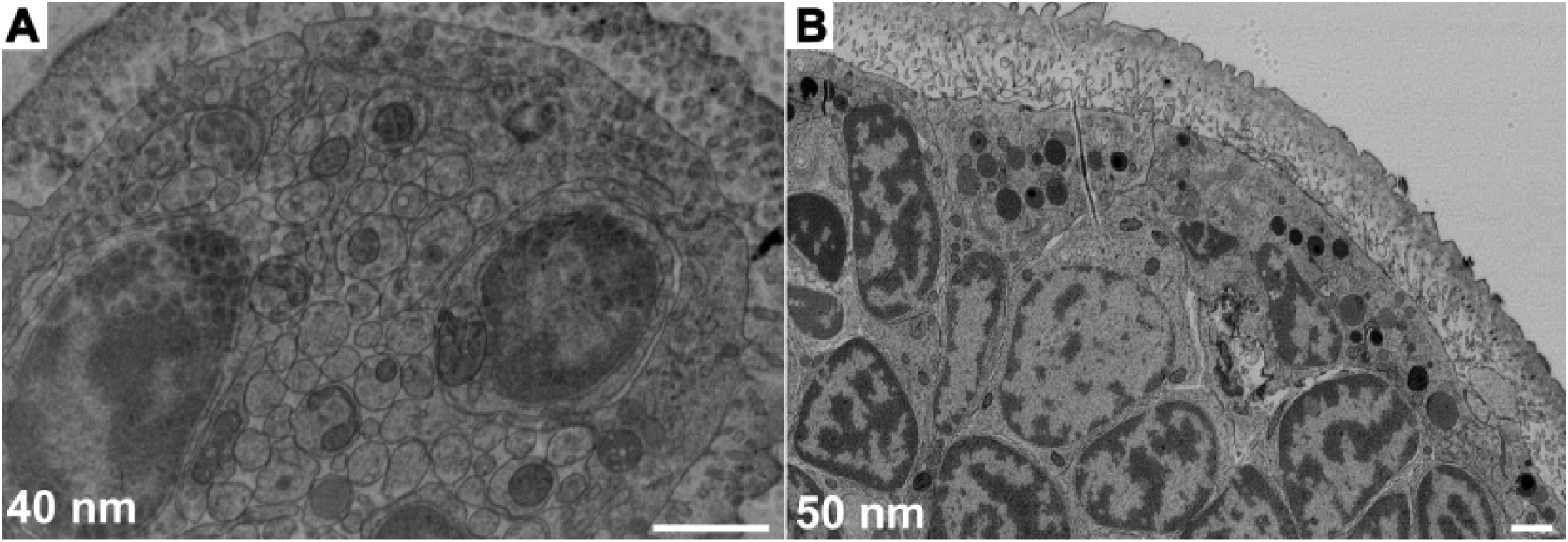
Sections of 6-day-old *Platynereis dumerilii* prepared with protocol C. (A) 40-nm-thick section, (B) 50-nm-thick section

We imaged post-stained and non-post-stained sections from samples from each of the different sample preparation protocols. Figure 7 shows a comparison between post-stained (B and D) and non-post-stained (A and C) samples. (A) and (B) are prepared following Protocol C, one of the best performing protocols, while (C) and (D) were prepared according to protocol D, a high-contrast protocol. In both cases, post-staining improved the contrast for AT.

**Figure 7:**
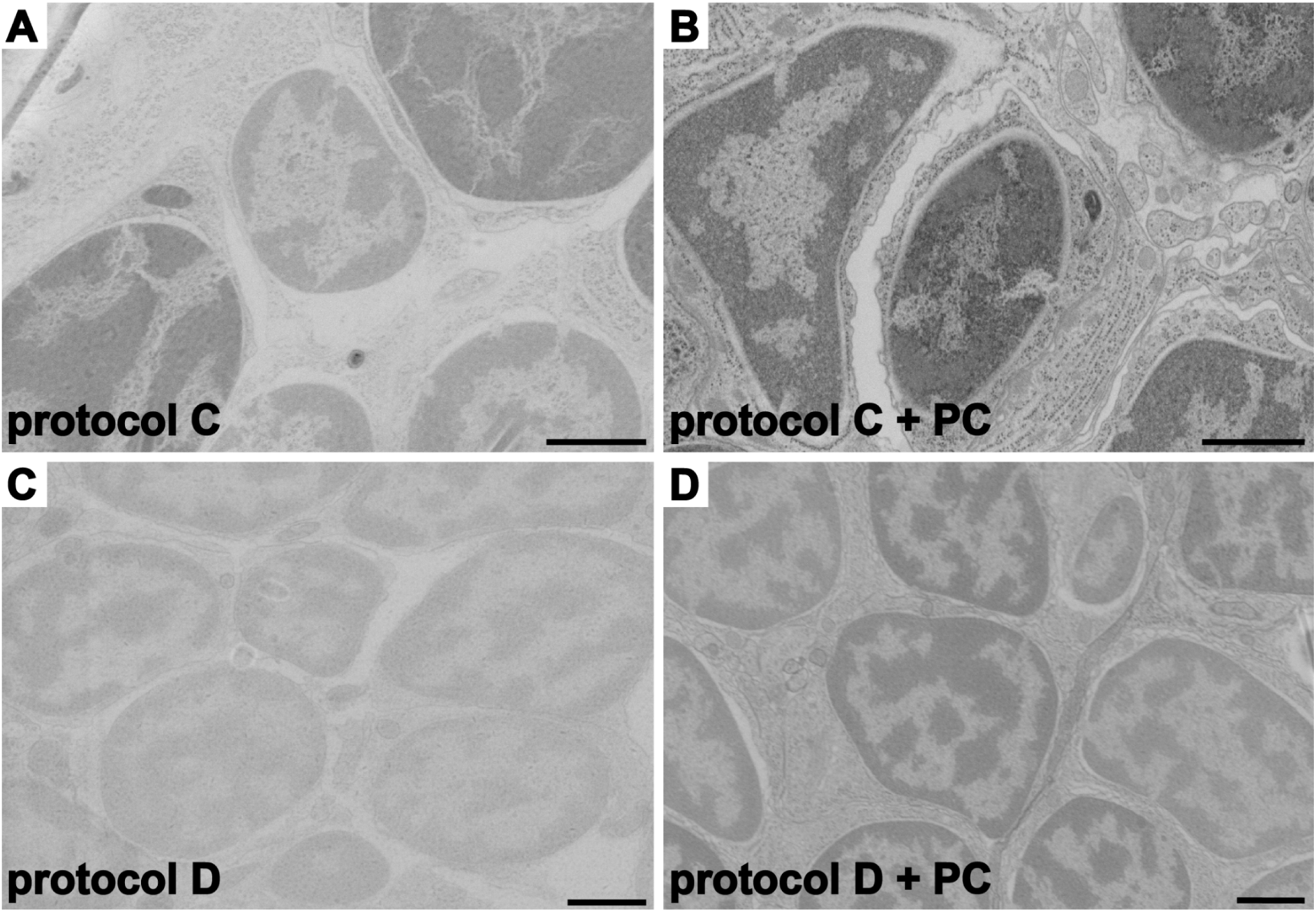
Comparison of post-stained and non-post-stained sections of different protocols. (A) SEM image of a non-post-stained section from a sample prepared with Protocol C, (B) image of a post-stained section of the same sample as (A). (C) SEM image of a non-post-stained section from a sample prepared with the high contrast Protocol D, (D) image of a post-stained section of the same sample as (C). Scale bars 1 μm.

We next assessed the SNR (Figure 3B) and contrast (Figure 3C) of the post-stained images after automatic brightness and contrast adjustment in Fiji. The SNR was highest for samples from protocol D (Figure 3B). The contrast values (Figure 3C) showed large variance even between samples of the same sample preparation.

While SNR and contrast are important variables, the most critical metric in EM is the quality of ultrastructural preservation. We therefore systematically compared preservation of cellular features by imaging various organelles (cilia, nuclei, golgi) across samples using similar imaging settings (Supplementary Figure 1).

The ultrastructure of the HPF-only samples was generally well preserved throughout different specimen and sample preparation protocols (Figure 4). Protocols A and C (Figure 4 and 4C) give good preservation and staining, especially Protocol C. These protocols still require on-section post-staining, which can introduce some variability.

Protocol B is a high-contrast protocol originally developed for brain serial-blockface EM that we selected in an attempt to obviate the need for post-staining. This protocol also gave good preservation and staining (Figure 4B), however post-staining was still required for optimal results. The lower stain uptake may be due to the reduced penetration of the staining agents through the collagenous cuticle of *Platynereis* larvae. After sectioning, we observed folds in all samples prepared with protocol B, suggesting that resin infiltration was incomplete.

Protocol D (Figure 4D) utilizes a large amount of heavy metals, which results in highly stained samples even without post-staining the sections. This results in high CNR and SNR values (Figure 3). However, at the ultrastructural level most cellular structures showed a similar level of contrast making it difficult to distinguish fine details. In addition, the membranes around the cilia appeared to be shrunk (Supplementary Figure 1).

Samples prepared with protocols E and F showed artefacts commonly associated with chemical fixation such as undulating membranes [27], [28], [29] (Figure 4E and F). Another consistent artefact that we observed in these samples was freeze damage in the nuclei (Figure 4E and F). We did not observe this phenomenon with HPF-only samples including samples processed in the same freezing session (Figure 4A-D).

We did not notice any major resin related artefacts, other than folds in all samples prepared with protocol B (Figure 4B).

## Discussion

The systematic vEM-based investigation of aquatic environmental samples and non-conventional marine organisms has recently become possible [12], [37], [38]. Aquatic and especially marine organisms can be challenging to prepare for volume electron microscopy [39], [38], [40]. Indeed, they can possess shells, a cuticle, mucus or other layers and structures, hampering contrasting or infiltration [39]. Osmolarity is also a critical factor with such samples, requiring the adaptation of buffers for chemical fixation [41], [42] or cryopreservation protocols. Our study aims to provide a set of optimised and reliable sample-preparation protocols for marine invertebrate larvae, tested here for the marine annelid *Platynereis dumerilii*. We also highlight the critical importance of using HPF over chemical fixation, previously leading to fixation artefacts [12].

Our results show that HPF followed by FS is the method of choice for vEM sample preparation of larval *Platynereis dumerilii*. We found that collecting 10-15 samples in one carrier and freezing them together gives samples that are optimal for X-ray imaging and further manipulation of the larvae. We also suggest carrying out freeze-substitution with samples in a freeze-substitution bucket to improve preservation by minimising sample handling during the most critical phase of the protocol. Using the buckets also allows a higher number of samples to be processed at the same time, 20 instead of 13 in the case of tubes, the number can be further increased to 40 and 18 respectively with custom inserts into the AFS2 [43]. Using the buckets also reduces the volume of solutions needed for the freeze-substitution and embedding. The use of freeze-substitution buckets has other advantages other than the slight increase in intact larva. For example the AFS2 machine can either hold 13 tubes (1 sample each) or 2 buckets (10 samples each), thus enabling the freeze-substitution of a larger number of carriers with buckets than with tubes. Another advantage of using the buckets is that the exchange of solutions happens through a central tube that connects to all the side tubes, where the carriers are. This makes a more gentle solution exchange possible, while also preventing the accidental aspiration and loss of specimens.

Analysis of CNR and SNR revealed differences between protocols. However, as we observed in our EM images, samples that have a higher SNR do not necessarily have better quality preservation. For example, protocol D had the highest CNR and SNR, but gave samples that were more homogeneously stained. Therefore the final choice of method should always rely on the expert evaluation of the EM images.

Overall, the best performing protocol on *Platynereis* larvae were protocols A and C. Our collaborators and we have also successfully used these protocols for other small invertebrates, including flatworm, priapulid and crustacean larvae (data not shown), suggesting that the protocols could be more widely used.

A limitation of this workflow is the intrinsic size-limits of HPF as a method for cryopreservation (around 200 µm) and the frequent occurrence of cracks, which likely appear during the freeze-substitution process (e.g., during transfer from -196 ℃ liquid nitrogen to -90 ℃). HPF also can not be used for animals with thick cuticles, as it can act as a thermal barrier [11]. Our results nevertheless show the advantages of cryopreservation (e.g., ultrastructure preservation, minimal artefacts) and we also propose strategies to minimise the drawbacks (e.g., cracks) of the workflow in order to obtain optimal results.

### Advantages of X-ray imaging in a volume EM workflow

Our work highlights the importance of the systematic screening of specimen integrity and contrast by high-throughput X-ray imaging. X-ray reconstructions also had the benefit of enabling the precise locating of the samples within the blocks. This information can be used to guide the trimming step. Precise trimming limits the sectioning to the necessary region of the sample, sparing the knife and also facilitating the start of sectioning [7]. The body posture of the specimens can also be determined based on the X-ray images. Animals that are for example bent (Figure 2B) or contracted can be identified and eliminated. This is not a necessary step, however having the specimens in a relaxed and neutral position can help to align them to images or atlases based on other modalities, such as light microscopy [30]. In addition, the X-ray images also enable the assessment of stain uptake of the sample and the identification of specimens with low amount of stain (Figure 2F) and high amount of stain (Figure 2G). We could also correlate X-ray image contrast to the contrast in EM images. Furthermore, the blocks that were prepared from the samples embedded in the bucket, contained several specimens. This made X-ray imaging a larger number of larvae faster and it also allowed us to identify the well preserved specimens that should be cut out and re-embedded.

## Conclusions

In this study, we introduce a sample preparation workflow for vEM, based on the detailed exploration of different protocols and parameter optimization at each step. This workflow is comprehensive and adaptable for different kinds of sensitive samples, and provides a useful tool for studying whole-organism morphology at subcellular resolution. The workflow allows the preservation of heterogeneous tissues, making it suitable for the study of cellular and subcellular structures of whole organisms and potentially whole-body connectomics. However, the protocol could also be suitable for other samples, such as cells, organoids or tissue samples (e.g., biopsies). By utilizing cryopreservation, the workflow we optimized here preserves the samples as close to their native state as possible therefore enabling downstream analyses at higher quality, potentially improving automated segmentation of subcellular elements [44].

## Acknowledgments

The authors would like to thank Dr. Réza Shahidi and Dr. Charlotta Fuyana from the Heidelberg University Electron Microscopy Core Facility for their continued support and expert training and advice. We are grateful to all colleagues who shared their in-house freeze-substitution protocols with us. We also thank Emily Savage for expertly managing our *Platynereis* facility and Elizabeth Duke and her team for support at the Hamburg P14 beamline. We acknowledge the European Synchrotron for the provision of beamtime under proposal number LS-3613, and we would like to thank Dr Peter Cloetens and the group of Dr Alexandra Pacureanu for assistance and support in using beamline ID16A. We would also like to thank the team of Alexandra Pacureanu for their support at the ESRF ID16A beamline. The authors also want to thank the two anonymous reviewers for their detailed and constructive feedback on a previous version of the manuscript. This work was supported by the European Union’s Horizon Europe research and innovation programme through the Marie Skłodowska-Curie Doctoral Network ZooCELL under Grant Agreement No. 101119891. This project received funding from the European Research Council (ERC) under the European Union’s Horizon 2020 research and innovation programme (grant agreement No. 101020792).

---

**Supplementary Figure 1.**
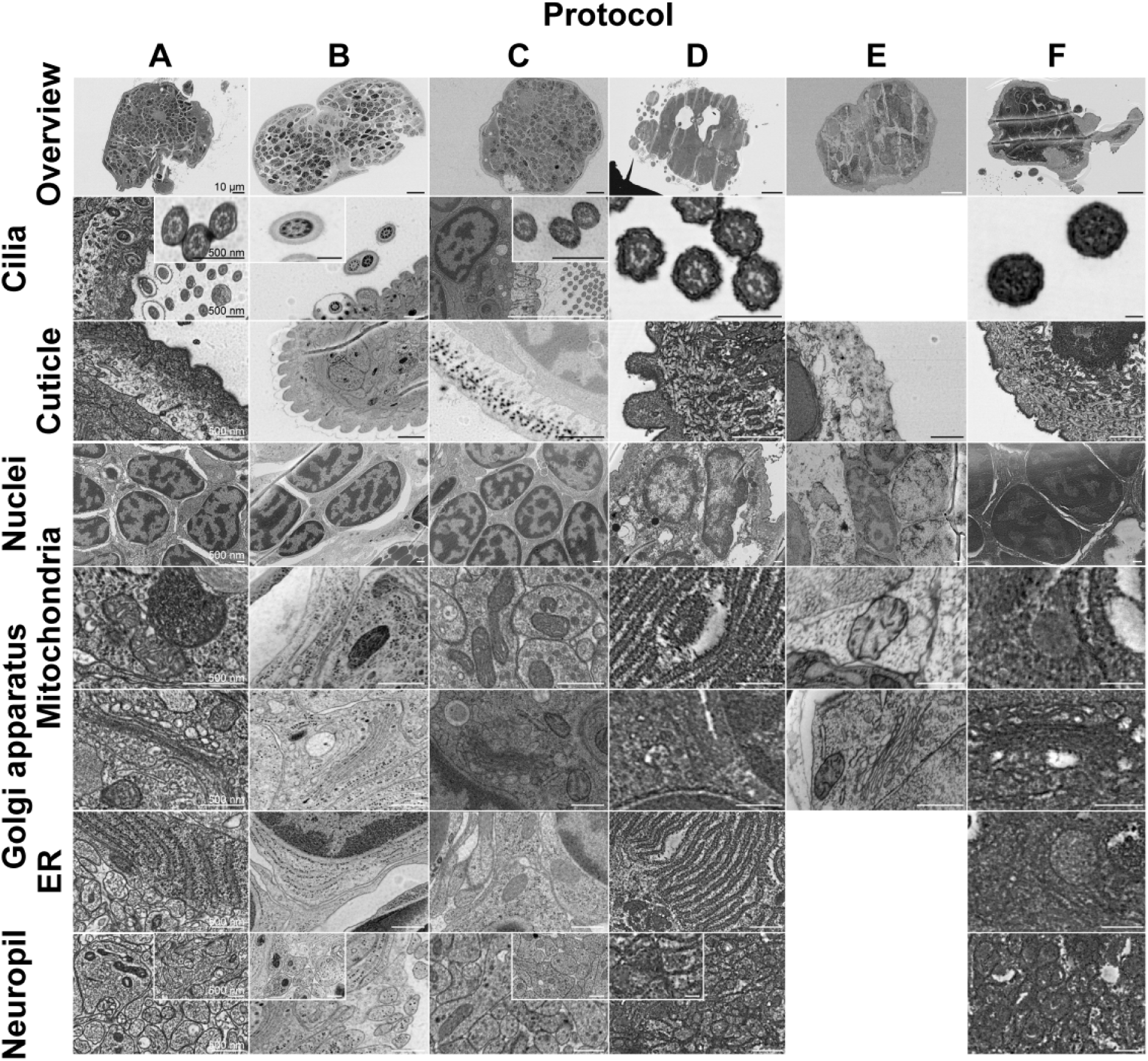
: Comparison of utrastructural preservation by SEM imaging between the different sample preparation protocols.

**Supplementary Figure 2.**
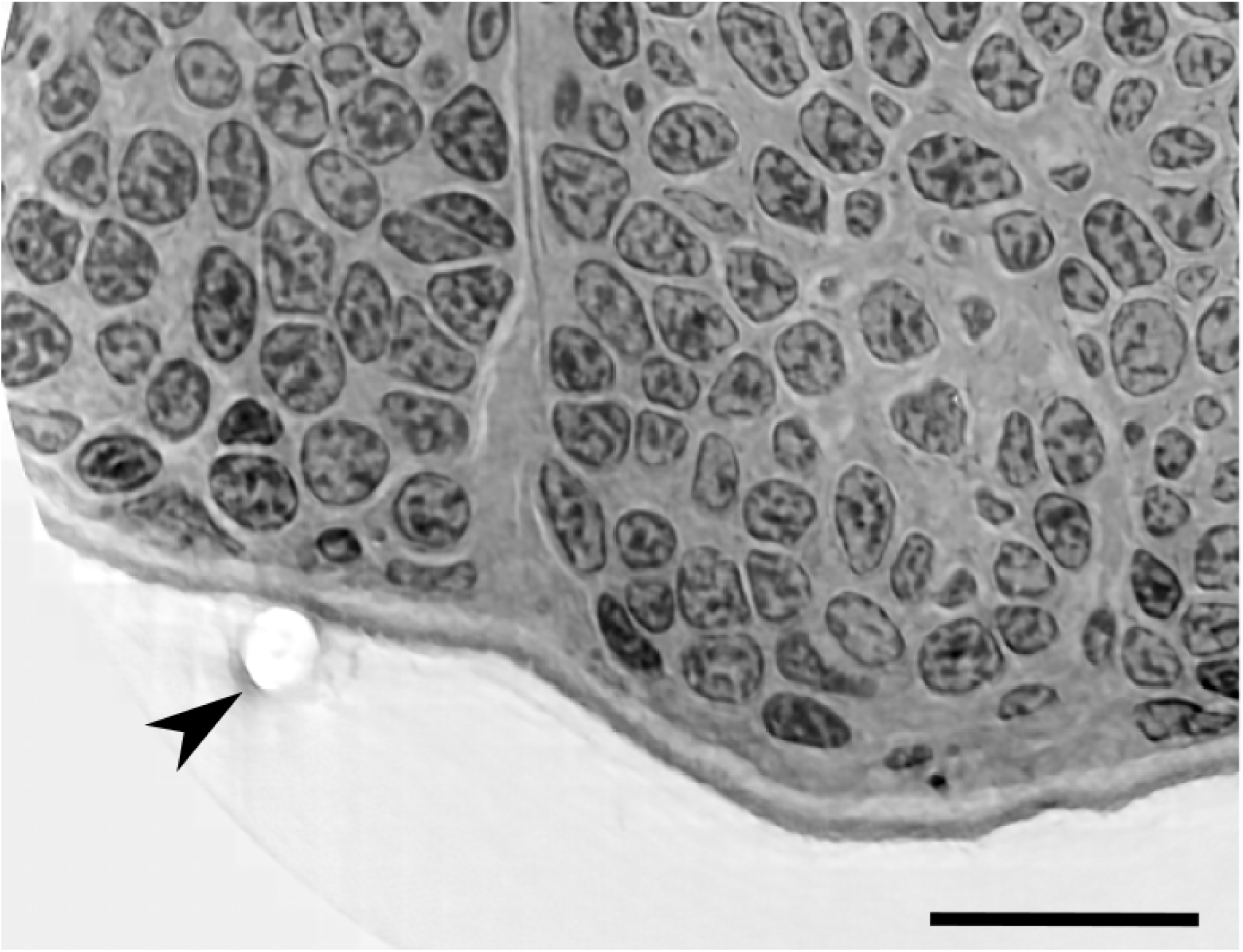
: Example of beam damage (arrowhead) of a *Platynereis dumerilii* sample scanned at 20 nm voxel size at ESRF. Scale bar 10 μm.

**Supplementary Table 1.** : Detailed information about reagents, consumables and equipment used in the experiments

Reagent, consumables and equipment table
| Chemicals | Full name | Company | Catalogue_number |
| --- | --- | --- | --- |
| 1-Hexadecene | - | Merk | 8.22964.0500 |
| OsO4 | Osmium tetroxide | EMS | 19134 |
| Acetone Glass distilled | - | Science Services | E10016 |
| LX112 | - | Ladd research | 21310 |
| NMA | Methyl-5-Norbornene-2,3-Dicarboxylic Anhydride | EMS | 19000 |
| DDSA | Dodecenyl Succinic Anhydride | EMS | 13710 |
| DMP-30 | Glycidether Accelerator DMP-30, | Carl Roth | 8621.1 |
| Tannic Acid | - | Thermo scientific | A17022.30 |
| Uranyl Acetate | - | Serva | 77870.01 |
| Glutaraldehyde | - | EMS | 16220 |
| MeOH | Methanol | Carl Roth | 5181.1 |
| HEPES |  | Merck | 54457-50G-F |
| Cacodylic acid | - | Serva | 15540-03 |
| Glycine | - | Fisher Scientific | G/0800/60 |
| Potassium ferrocyanide | - | Omikron | 90-0447 |
| Potassium ferrocyanide | - | Merck | P9387 |
| CaCl<sub>2</sub> | Calcium Chloride | Merck | C5670 |
| TCH | Thiocarbohydrazide | EMS | 21900 |
| EtOH | Ethanol | Merck | 32221-1L-M |
| PFA | Paraformaldehyde | EMS | 15710 |
| NaCl | Sodium chloride | Carl Roth | 0601.1 |
| Na<sub>2</sub>HPO<sub>4</sub>*2H<sub>2</sub>O | Di-sodium hydrogen phosphate dihydrate | Carl Roth | T877.1 |
| KH<sub>2</sub>PO<sub>4</sub> | Potassium dihydrogen phosphate | Carl Roth | 23Y6.2 |
| Lead citrate | - | Science Services | DM22410 |
| **Consumables** | **Full name** | **Company** | **Catalogue_number** |
| Reagent bath | - | Leica microsystems | 16707154 |
| Flow through rings | - | Leica microsystems | 16707157 |
| Cryotubes | - | Heathrow scientific | HS10060 |
| Carrier A gold/copper 0.2 -0.1 mm | - | Wohlwend GmbH | 662 |
| Carrier A Aluminium 0.2 -0.1 mm | - | Wohlwend GmbH | 241 |
| Carrier B Aluminium 0.3 mm | - | Wohlwend GmbH | 242 |
| ITO glass | Indium-tin-oxide coated glass slides | UQG (Optics) Ltd. | CIO-2258 |
| **Equipment** | **Full name** | **Company** |  |
| AFS2 | - | Leica microsystems |  |
| PELCO BioWave Pro+ | - | Pelco |  |
| HPM10 | - | Baltech |  |
| GeminiSEM 500 | - | Zeiss |  |
| GeminiSEM 360 | - | Zeiss |  |
| UC7 Ultramicrotome | - | Leica microsystems |  |
| Glow discharger easiGlow | - | PELCO |  |

**Supplementary Table 2.** : Detailed description of protocol A in 6-day-old *Platynereis dumerilii*

| temperature start | temperature end | duration | slope | other comments |
| --- | --- | --- | --- | --- |
| -90 | -90 | 37:00:00 | 0°C/h | 1% OsO<sub>4</sub>, 0.5% UA, 3% water in dry acetone |
| -90 | -60 | 15:00 | 2°C/h | 1% OsO<sub>4</sub>, 0.5% UA, 3% water in dry acetone |
| -60 | -60 | 12:00 | 0°C/h | 1% OsO<sub>4</sub>, 0.5% UA, 3% water in dry acetone |
| -60 | -30 | 15:00 | 2°C/h | 1% OsO<sub>4</sub>, 0.5% UA, 3% water in dry acetone |
| -30 | -30 | 12:00 | 0°C/h | 1% OsO<sub>4</sub>, 0.5% UA, 3% water in dry acetone |
| -30 | 0 | 01:00 | 30°C/h | 1% OsO<sub>4</sub>, 0.5% UA, 3% water in dry acetone |
| 0 | 0 | 01:00 | 0°C/h | 1% OsO<sub>4</sub>, 0.5% UA, 3% water in dry acetone |
| 0 | -30 | 01:00 | 30°C/h | 1% OsO<sub>4</sub>, 0.5% UA, 3% water in dry acetone |
| -30 | -30 | 01:30 | 0°C/h | Acetone rinses |
| -30 | -10 | 02:00 | 10°C/h | 30% resin |
| -10 | 10 | 02:00 | 10°C/h | 50% resin |
| 10 | 20 | 02:00 | 5°C/h | 70% resin |
| 20 | 20 | overnight | 0°C/h | 100% resin |
| 20 | 20 | 03:00 | 0°C/h | 100% resin |
| 20 | 20 | 03:00 | 0°C/h | 100% resin |
| 20 | 20 | 03:00 | 0°C/h | 100% resin |
| 20 | 20 | overnight | 0°C/h | 100% resin + hardener |
| 20 | 20 | 08:00 | 0°C/h | 100% resin + hardener |
| 60 |  | 48:00:00 |  | curation |

**Supplementary Table 3.** : Detailed description of protocol B in 6-day-old *Platynereis dumerilii*

| temperature start | temperature end | duration | slope | other comments |
| --- | --- | --- | --- | --- |
| -90 | -90 | 54:00:00 | 0°C/h | FS1=2%OsO4, 0.5% UA, 1% GA in acetone |
| -90 | -20 | 08:00:00 | 8.75°C/h | FS1=2%OsO4, 0.5% UA, 1% GA in acetone |
| -20 | -20 | 12:00:00 | 0°C/h | FS1=2%OsO4, 0.5% UA, 1% GA in acetone |
| -20 | 4 | 02:24:00 | 10°C/h | FS1=2%OsO4, 0.5% UA, 1% GA in acetone |
| 4 | 4 | 00:40:00 | 0°C/h | acetone washes |
| 4 | 4 | 14:30:00 | 0°C/h | FS2=1% UA in acetone |
| 4 | 20 | 00:30:00 | 32°C/h | FS2=1% UA in acetone |
| 20 | 20 | 00:40 | 0°C/h | anhydrous acetone washes |
| 20 | 20 | 02:00 | 0°C/h | 25% resin in acetone |
| 20 | 20 | 02:00 | 0°C/h | 50% resin in acetone |
| 20 | 20 | 02:00 | 0°C/h | 75% resin |
| 20 | 20 | overnight | 0°C/h | 100% resin |
| 20 | 20 | 48:00:00 | 0°C/h | 100% resin |
| 20 | 20 | 04:00 | 0°C/h | 100% resin with DMP30 |
| 20 | 20 | 04:00 | 0°C/h | 100% resin with DMP30 |
| 20 | 20 | overnight | 0°C/h | 100% resin with DMP30 |
|  | 60 | 48:00:00 |  | oven |
|  | - | ready |  |  |

**Supplementary Table 4.** : Detailed description of protocol C in 6-day-old *Platynereis dumerilii*

| *temperature start* | *temperature end* | *duration* | *slope* | *other comments* |
| --- | --- | --- | --- | --- |
| -90°C | -90°C | 28:00:00 | 0°C/h | 0.1% TA and 0.5% GA in acetone |
| -90°C | -90°C | 01:30 | 0°C/h | Acetone washes |
| -90°C | -90°C | 29:30:00 | 0°C/h | 2% OsO4 in acetone |
| -90°C | -20°C | 14:00 | 5°C/h | 2% OsO4 in acetone |
| -20°C | -20°C | 16:00 | 0°C/h | 2% OsO4 in acetone |
| -20°C | 4°C | 04:00 | 6°C/h | 2% OsO4 in acetone |
| 4°C | 4°C | 01:30 | 0°C/h | Acetone washes |
| 4°C | 20°C | 02:00 | 8°C/h | 25% resin in acetone |
| 20°C | 20°C | 02:00 | 0°C/h | 50% resin in acetone |
| 20°C | 20°C | 02:00 | 0°C/h | 75% resin |
| 20°C | 20°C | overnight | 0°C/h | 100% resin |
| 20°C | 20°C | 48:00:00 | 0°C/h | 100% resin |
| 20°C | 20°C | 04:00 | 0°C/h | 100% resin with DMP30 |
| 20°C | 20°C | 04:00 | 0°C/h | 100% resin with DMP30 |
| 20°C | 20°C | overnight | 0°C/h | 100% resin with DMP30 |
|  | 60°C | 48:00:00 |  | oven |

**Supplementary Table 5.**
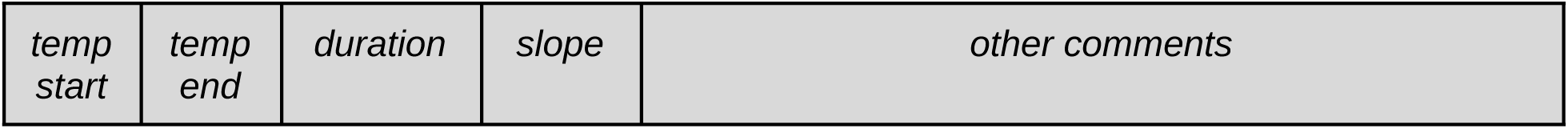
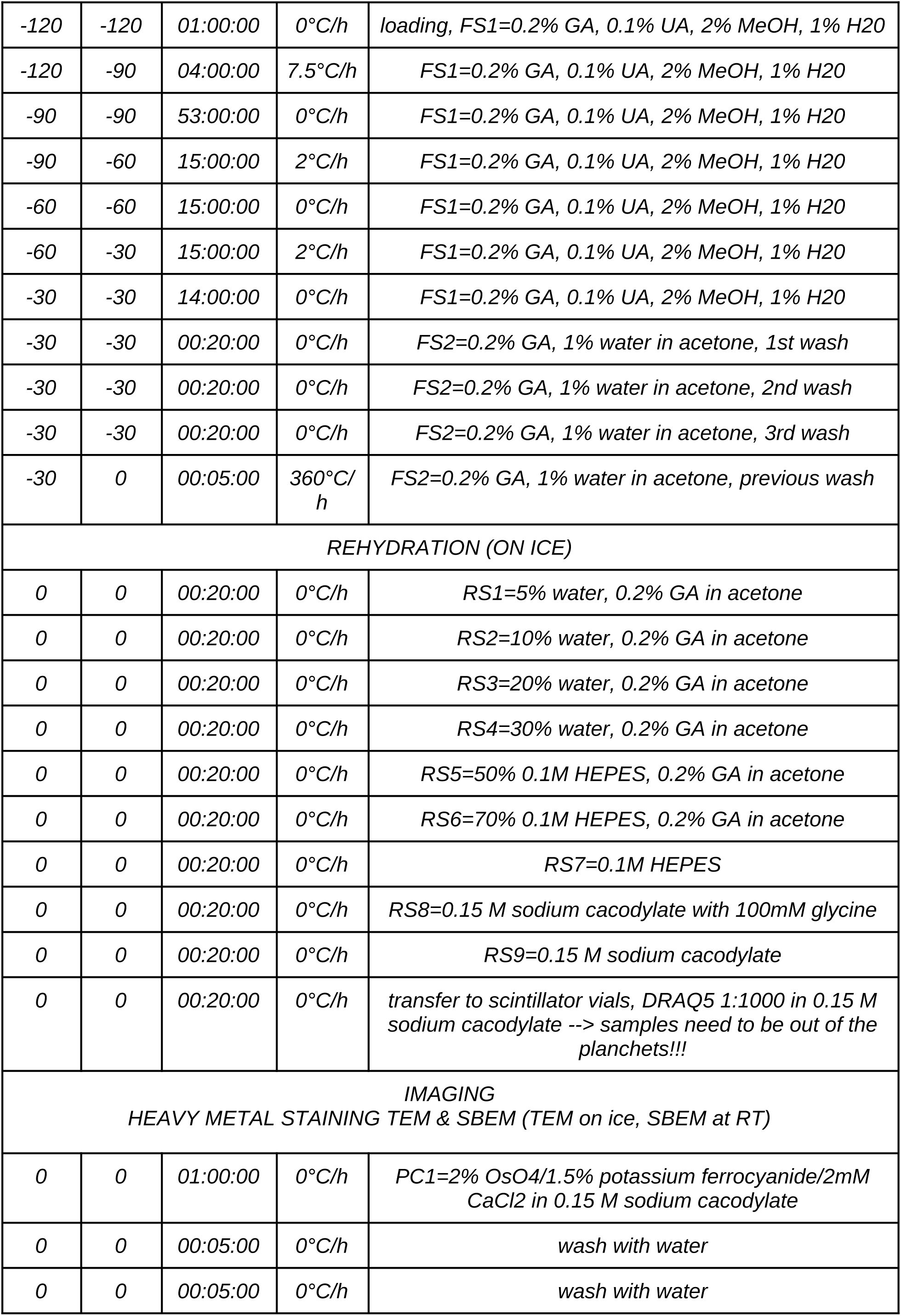
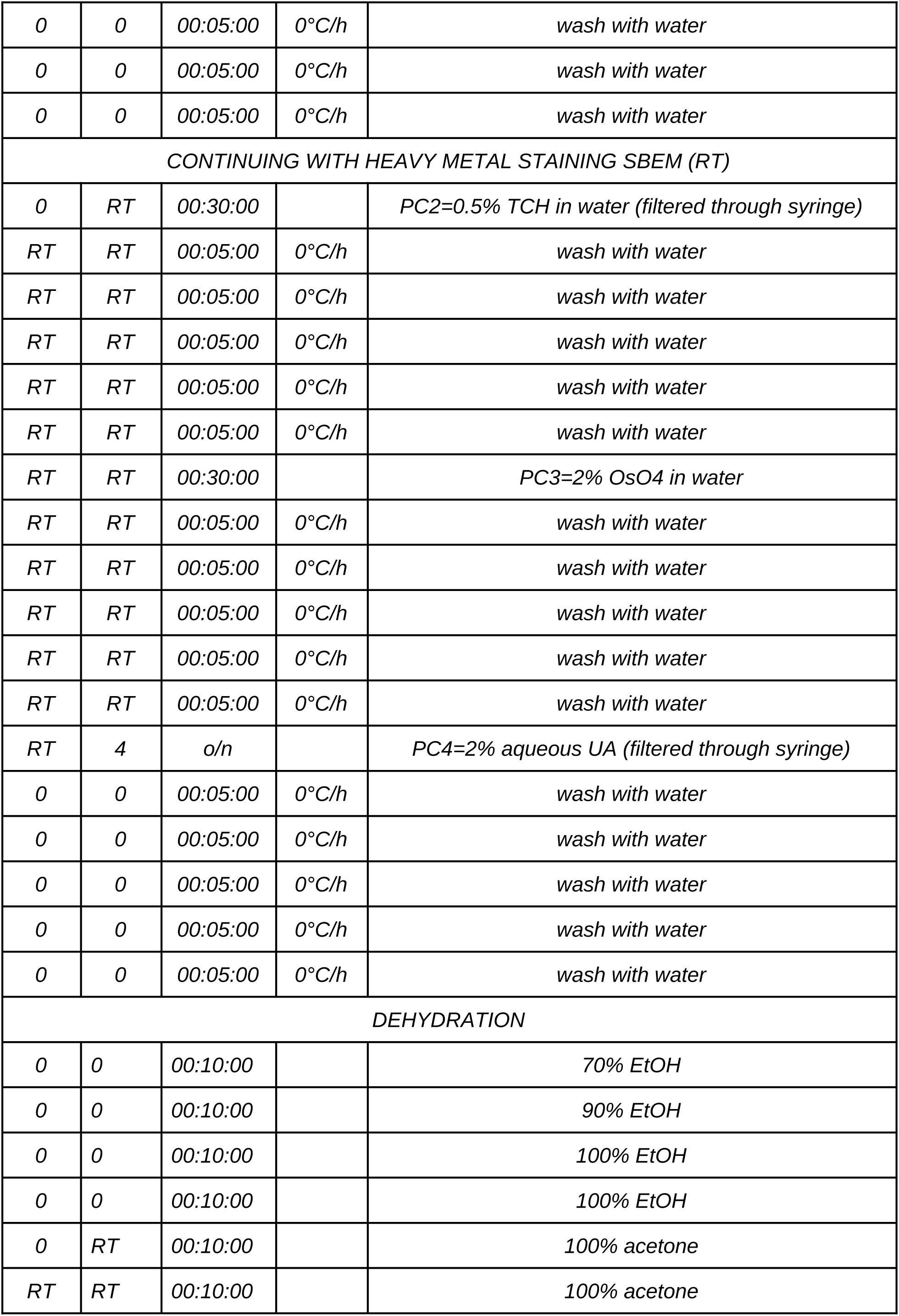
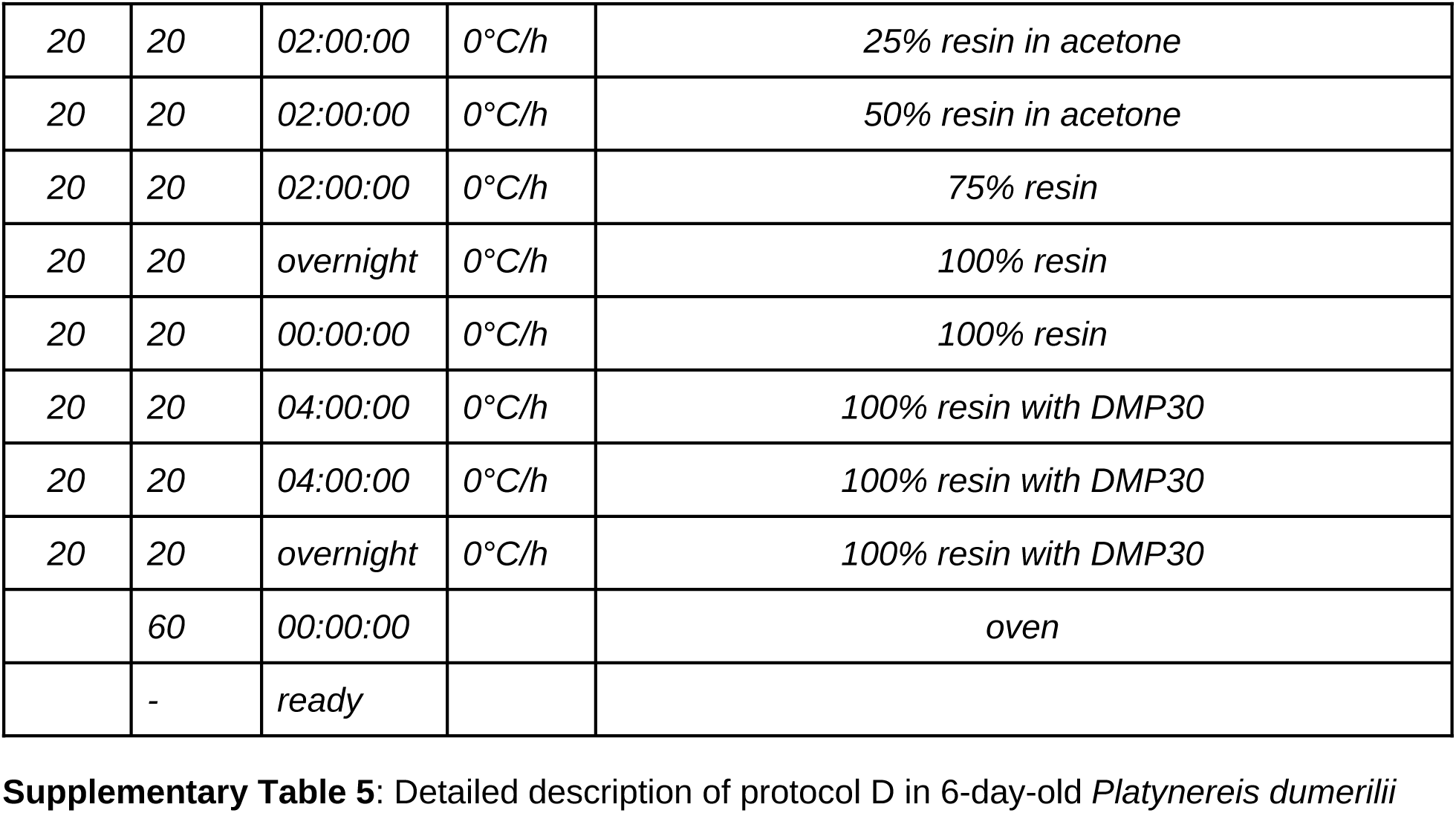
: Detailed description of protocol D in 6-day-old *Platynereis dumerilii*

